# Genome Wide Association Study in the New Haven Lexinome Project Identifies *GARRE1* as a Novel Gene for Reading Performance

**DOI:** 10.1101/2021.01.05.423827

**Authors:** Andrew K. Adams, Emily L. Guertin, Dongnhu T. Truong, Elizabeth G. Atkinson, Mellissa M.C. DeMille, Joan M. Bosson-Heenan, Jan C. Frijters, Jeffrey R. Gruen

**Affiliations:** Department of Pediatrics, Yale School of Medicine, New Haven, CT 06519; Department of Child and Youth Studies, Brock University, St. Catharines, ON L2S 3A1; Analytic and Translational Genetics Unit, Massachusetts General Hospital, Boston, MA 02114; Department of Genetics, Yale School of Medicine, New Haven, CT 06519; Investigative Medicine Program and Computational Biology and Bioinformatics, Yale School of Medicine, New Haven, CT 06519

## Abstract

Despite high prevalence and high heritability, few candidate genes have been identified for reading disability. To identify novel genetic variants we performed a genome-wide association study (GWAS) using high-depth whole genome sequencing and predicated on reading performance in 407 subjects enrolled in a longitudinal study of response-to-intervention, called the New Haven Lexinome Project. The primary GWAS identified a single peak of 31 SNPs on chromosome 19 that achieved the threshold for genome-wide significance (rs2599553 *P*=3.13×10^−8^) located over an expression quantitative trait locus (eQTL) for *GARRE1* (Granule Associated Rac And RHOG Effector 1). Little is known about the function of *GARRE1*, except that it is highly and developmentally expressed in human cerebellum relative to cortex. Local ancestry regression showed the strongest association for the lead variant in African or Admixed American populations, who have been under-represented in previous genetic studies of reading. We replicated our chromosome 19 results in the Genes, Reading, and Dyslexia (GRaD) cohort and found a moderating effect of age with implications for the consideration of developmental effects in the design of future analyses. Growth curve modeling demonstrated that minor alleles of the lead SNP are related to reading longitudinally from Grade 1 to Grade 5, and that children with at least 1 minor allele of rs2599553 persistently underperformed relative to their peers by 0.33 to 0.5 standard deviations on standardized assessments of non-word decoding and reading fluency.

**Significance Statement:** To the best of our knowledge, this work represents the only GWAS predicated on longitudinal reading performance data. Starting with initial discovery, we replicate our association in a second cohort, address common causes of type I error, localize the signal to a single gene, implicate a region of the brain most likely to be affected by variation in our candidate, show a gene-by-age effect with implications for study design in this field, and demonstrate that minor alleles of our lead SNP are associated with significant and persistent clinical effects on reading development in children.

## Introduction

Reading disability (RD) is a lifelong disorder beginning in childhood with significant long-term consequences including limited educational attainment and earning potential (Schatschneider and Torgesen, 2004; Peterson and Pennington, 2015). Despite advances in remediation methods (Wanzek et al., 2010, 2013), a significant barrier to effective intervention is early identification, and within school systems formal diagnoses are often delayed until the third grade (Lyon, et al., 1997). The earlier that reading problems are identified, the more likely intervention will be effective in remediating reading skill (Lovett et al., 2017). Reading performance is a complex function of language exposure, genetic, and developmental factors, making it difficult to distinguish children with RD from otherwise low performing peers (Snowling et al., 2003). Distinguishing between children with low initial ability who will have a persistent learning disability from those who respond well to regular instruction is critical for tailoring interventions and allocating school resources (Dion et al., 2010).

One promising approach to early identification is through screening for genetic variants associated with RD. Reading performance has a large genetic component with heritability estimates ranging from 30% to 70% (Byrne et al., 2002; Gayán and Olson, 2003; Harlaar et al., 2005). Genetic variants could also help to tailor interventions to specific needs of the child – critical components of a successful precision education program. Similar to precision medicine, this is a nascent specialty with the goal of developing interventions informed by variation in specific reading strengths and deficits, tailored to individual needs, and delivered within the optimal developmental window. The first step in building a successful precision education program is to identify informative genes and variants. However, studies with the aim to identify new genes associated with reading are generally underpowered, and identify variants with small effect sizes. Most genes associated with reading were discovered through family-based methods that identify only rare variants or candidate gene approaches from suspected pleiotropic traits (i.e. autism, educational attainment, specific language impairment, etc.). Few genes and variants have been identified through GWAS (Eicher et al., 2013; Gialluisi et al., 2014, 2019; Truong et al., 2019).

In addition, diverse and minority populations are chronically understudied. With the exception of a multivariate GWAS predicated on rapid automatized naming tasks in a cohort of adolescents of African and Hispanic ancestry (Truong et al., 2019), no other GWAS of reading or component skills in minority populations has been published to date. Understanding the full spectrum of the genetics of reading performance requires the inclusion of diverse populations (Sirugo et al., 2019).

For the current study, the primary GWAS and growth curve analyses were performed on data from the New Haven Lexinome Project (NHLP). The NHLP is a longitudinal study of the genetics of reading performance for children in Grades 1 through 5 from a diverse, urban, public school district. It was designed to address two prerequisites of a successful precision education program — early identification of RD risk and identifying predictors of intervention response. Every subject had serial assessments from psychoeducational batteries, extensive documentation of the home language environment, family history of reading problems and medical history, and high-depth whole genome sequencing. Assessments of academic skills included single word reading, phonological decoding, passage comprehension, math performance, attention and motivation.

Below we present the results of a novel GWAS predicated on single-word decoding performance, a critical component of reading fluency, which identified an association between reading and the gene *GARRE1*. *GARRE1* is differentially expressed in the cortex and cerebellum postnatally, in support of neuroimaging associations between the cerebellum and reading ability (Miller et al., 2014). We replicated our results in the cross-sectional Genes Reading and Dyslexia (GRaD) Study, comprised of Hispanic American and African American youth, but drawn from multiple regions within the US. Finally, we investigated the gene-by-development effect of age in the GRaD cohort and the effect of minor alleles from our locus peak on the development of reading skill through growth curve analysis.

## Methods

### The New Haven Lexinome Project

Data were analyzed from 420 children enrolled in Grade 1 classes across 31 public elementary schools located in New Haven, CT in 2015 and 2016 (Table 1). Prospective parents and their children were invited to participate by a letter from the Assistant Superintendent and Director of Research at New Haven Public Schools. Parents or guardians who expressed interest in participating were contacted by a study recruiter who explained the study in English or Spanish and obtained informed consent from the prospective parent or guardian, and verbal (under the age of 7) or written assent from the child. Exclusion criteria were history of a developmental delay, pervasive developmental disorder, autism spectrum disorder, neurological disorder or epilepsy, brain injury, gestational age less than 37 weeks, prolonged hospitalization in a neonatal intensive care unit, or greater than 20 days of unexcused absences in the prior school year. All children provided a saliva sample for subsequent DNA extraction (DNA Genotek, Oragene, Kanata, Ontario, Canada). This study was approved by and receives on-going review from the Human Research Protection Program at Yale University (IRB #1507016121).

**Table 1.** NHLP GWAS sample demographics (N = 407 due to sparsely missing covariate data).

|  |  |
| --- | --- |
| Sex (M : F) | 192 : 215 |
| Age (Years; mean) | 6.0 - 10.25 (7.42) |
| SES (High : Low) | 43 : 364 |
| Mean-split Decoding Composite (Below mean : Above mean) | 179 : 228 |
| Self-Report European Ancestry (N subj.) | 19 |
| Self-Report African Ancestry (N subj.) | 95 |
| Self-Report Hispanic Ancestry (N subj.) | 107 |
| Self-Report Dual Ancestry (N subj.) | 99 |
| Failed to Report Ancestry (N subj.) | 96 |

Participants in the NHLP were evaluated longitudinally with a standard battery of reading and psychoeducational assessments twice per academic year from the start of Grade 1 through the start of Grade 5. The reading battery included the sight word efficiency and phonetic decoding efficiency subtests of the Test of Word Reading Efficiency 2^nd^ edition (TOWRE-2), seven subtests from the Woodcock-Johnson III Tests of Achievement 3^rd^ edition (WJ-III), seven subtests from the Comprehensive Test of Phonological Processing 2^nd^ edition (CTOPP-2), the Gray Oral Reading Tests 5^th^ edition (GORT), the Morphological Awareness Task, and the Lexinome Reading Motivation Measure (Wagner et al., 2012.; Braden and Alfonso, 2003; Torgesen, et al., 2012; Wiederholt and Bryant, 2012). The psychoeducational battery included the Clinical Evaluation of Language Fundamentals Screening Test (CELF-5 5^th^ edition), Peabody Picture Vocabulary Test (PPVT 4^th^ edition), Wechsler Abbreviated Scale of Intelligence (WASI-II 2^nd^ edition), A Developmental NEuroPSychological Assessment (NEPSY-II 2^nd^ edition), the Wechsler Intelligence Scale for Children (WISC-IV 4^th^ edition), the Barkley ADHD Screening Checklist Rating Scale (parent), Strengths and Weaknesses in ADHD-Symptoms and Normal Behavior (SWAN parent), Sluggish Cognitive Tempo Scale (parent), Musical Meter Differentiation, Rhythm Entrainment Task, and Stork Task (Wechsler, 1949, 2011; Dunn and Dunn, 2007; Brooks et al., 2009; Barkley, 2011; Jacobson et al., 2012; Swanson et al., 2012; Wiig et al., 2013)

### DNA Collection and Sequencing

DNA was extracted from saliva samples and sequenced at the Yale Center for Genome Analysis (YCGA). Saliva samples with high levels of bacterial contamination or evidence of poor quality were excluded and recollection was attempted. DNA samples that met all QC requirements were sequenced on one of two platforms, the Hi-Seq 2500 or the Novaseq (Illumina, Inc. San Diego, CA). DNA samples were sequenced to a mean depth of 30x to ensure that all variants were captured.

Sequence reads were aligned to the hg19 reference genome using the memory efficient algorithm in the Burrows-Wheeler Aligner (bwa-mem) to generate binary compressed aligned sequences for each subject (Li and Durbin, 2009). Variants from reference hg19 were identified using the Genome Analysis Toolkit (GATK; v3.7) according to best practices published by the Broad Institute (McKenna et al., 2010; Van der Auwera et al., 2013). Duplicate reads arising from a single DNA segment were marked, and BAM quality scores were recalibrated before variant calling. Variants were called using the HaplotypeCaller to generate a single genomic variant call file (gVCF) for each subject. Subject-level gVCFs were broken into 1megabasepair (Mbp) segments and imported into GenomicsDBImport (GATK v.4). Joint called segment gVCFs were combined into a single GVCF for all 420 subjects. The GenotypeGVCFs module was used to convert the full cohort GVCF into a single VCF containing all variants across all subjects.

This all-variants VCF was subjected to Variant Quality Score Recalibration (VQSR) according to Broad Institute best practices.(DePristo et al., 2011) To train the model for SNPs, HapMap and the Omni array data provided by the Broad were treated as truth sets, and the 1000 Genomes Project was used as a resource for both true and false variants. (Gibbs et al., 2003; The Genomes Project Consortium, 2015) dbSNP was not used to train the model, but was used to evaluate transition/transversion (Ti/Tv) ratios.(Sherry et al., 1999) To detect insertions and deletions (indels), the Mills gold standard resource provided by the Broad was used as a model-training truth set.(Mills et al., 2011) Variants were scored either PASS or into three additional tranches of lower confidence. All variants that did not get assigned PASS by the VQSR model were filtered out of further analyses.

The output VCF containing all variants for the NHLP cohort was filtered for a minimum variant sequencing depth of at least 10x and indels were excluded from further study. This final SNP-only VCF was converted to binary PLINK format for association analysis. Subjects were updated with self-declared sex codes derived from the subject questionnaire. Sex assignment was cross-checked based on X-chromosome genotypes (no discrepancies were observed). X-chromosome, Y-chromosome and mitochondrial sequences were excluded following sex check. Remaining variants were filtered for minor allele frequency (MAF) > 5%, variants with cohort missingness above 5% were removed, and variants with Hardy-Weinberg equilibrium p-values below 0.00001 were excluded. The remaining 6,278,349 variants were included in downstream analyses.

The final sample was evaluated for cryptic relatedness using the KING v2.2.4 algorithm.(Manichaikul et al., 2010) Using the cleaned and quality controlled set of SNPs, relationships between subjects were estimated using the ‘--related’ option. Briefly, all pairwise relationship coefficients were calculated and compared to expected values based on pedigree status, under the assumption that all subjects were unrelated. Subjects with relatedness above expected were flagged for follow-up examination. Through this procedure we detected one pair of siblings and one subject from this pair was excluded from all analyses.

### Latent Reading Phenotype

A latent reading score was generated for each subject using covariance modeling performed on the following tasks (Table 2): WJ-III Word Attack (nonword decoding), WJ-III Word Identification (real word decoding), TOWRE Sight Word Efficiency (timed real word decoding), TOWRE Nonword Decoding (timed non-word decoding), and GORT Accuracy (real word decoding in context) to generate a single, normally distributed distillation of decoding performance for each subject. Missing testing data were backfilled from earlier or later testing dates to maximize sample size, though all testing used was pre-intervention to prevent confounding.

**Table 2.** Measures performed in the NHLP. Measures included in the decoding composite phenotype are in bold.

| Measure | Subtests | Targeted Construct(s) | Author |
| --- | --- | --- | --- |
| **Test of Word Reading Efficiency, 2nd Ed. (TOWRE-2)** | **-Sight Word Efficiency**<br>**-Phonetic Decoding Efficiency** | Reading accuracy and fluency for whole words and nonwords | Torgesen, Wagner, & Rashotte, 2012 |
| **Woodcock-Johnson III Tests of Achievement (WJ III ACH)** | **-Letter-Word ID**<br>-Reading Fluency<br>-Passage Comprehension<br>**-Word Attack**<br>-Calculation<br>-Math Fluency<br>-Applied Problems | Academic achievement in a variety of reading and math tasks targeting specific cognitive abilities | Woodcock, McGrew, & Mather, 2001 |
| **Gray Oral Reading Tests, 5th Ed. (GORT-5)** | **-Reading Fluency**<br>-Reading Comprehension | Oral reading ability including rate, accuracy, fluency and comprehension | Wiederholt & Bryant, 2012 |

Latent reading composite scores were converted to Z-scores using R (v3.6.1), which were used to assign a binary status for downstream analysis (R Core Team, 2016). Subjects with Z-scores below the mean were assigned pseudocase status. Subjects with Z-scores above the mean were assigned pseudocontrol status (Figure 1). The mean standard score reading performance for the pseudocases was 86.7 (SD = 11.5), and was significantly lower (*t*(475) = 19.1, p = 6 x 10^−6^) than the pseudocontrols at 105.5 (SD = 9.9). The roughly equivalent numbers of pseudocases (47% of *N* = 415) and pseudocontrols reflects the intentional enrichment for subjects at risk for reading failure that was inherent in the design of the NHLP.

**Figure 1.**
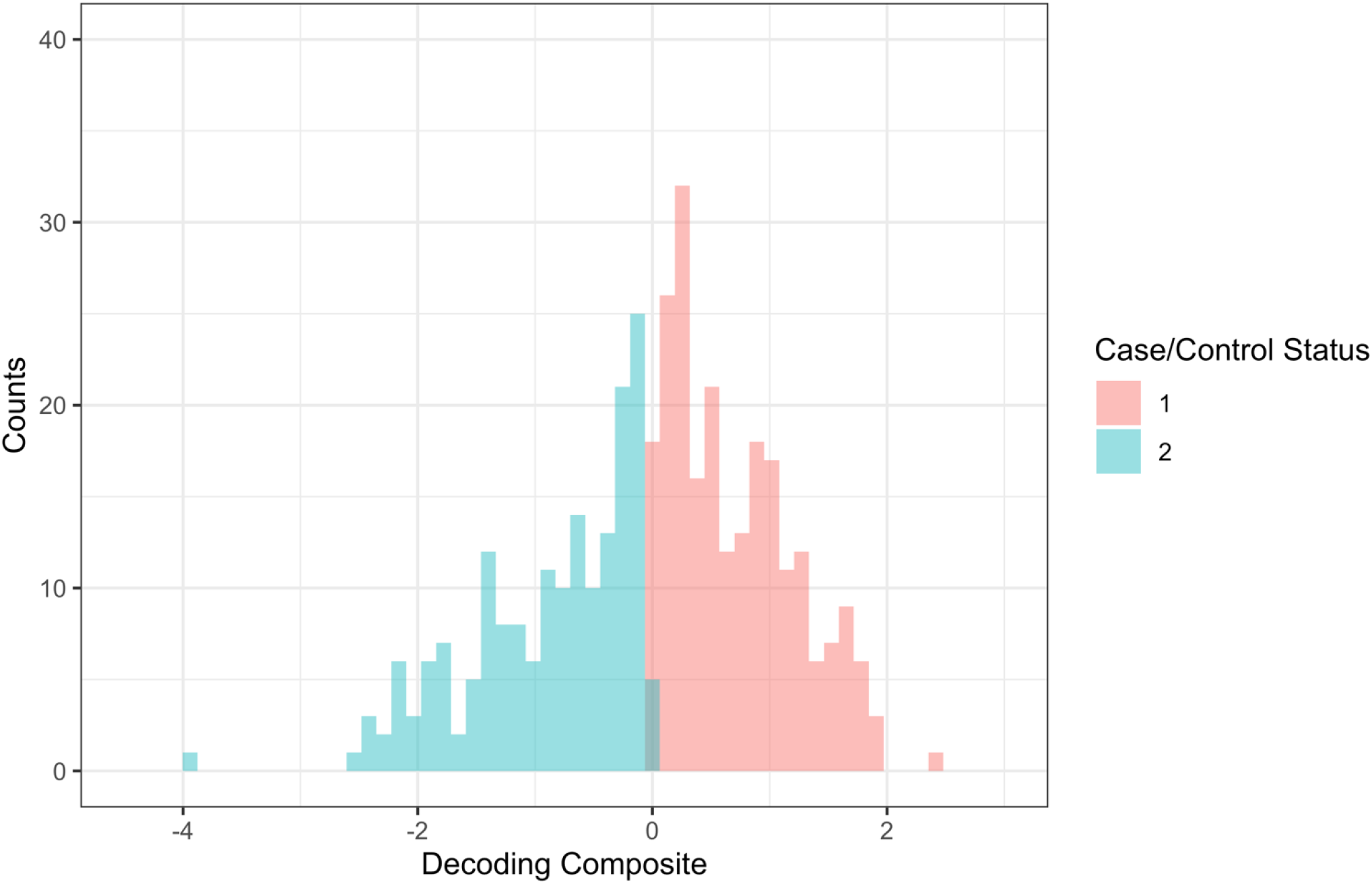
Case/control dichotomization of the decoding composite (N = 415) in the NHLP.

### Socioeconomic Status Assignment

Socioeconomic status (SES) was included as a covariate in all the analyses using a binary classification as previously described (Truong et al. 2019). Subjects who reported participation in at least one government assistance program were assigned to the low SES category. Subjects that did not report participation in any government assistance program, had a reported income that would have made them ineligible to receive government assistance, or left this questionnaire item incomplete, were assigned to the average or above SES category. Subjects with missing SES self-report or missing income data that were also recorded as receiving free or reduced price lunch in data provided by New Haven Public Schools, were assigned to the low SES category. If SES could not be determined, subjects were assigned to the low SES category (58 subjects), since 86% of families who responded to the SES questions received some level of government assistance.

### Additional NHLP Covariates

Age at the time of testing was used for all the analyses to construct the latent reading score, and as a covariate in all models. For subjects with backfilled pre-intervention testing data, age was assigned when the majority of testing was performed. Sex was included as a covariate in all models.

### Ancestry Analysis

A principal components analysis was used to correct for population stratification and to control for admixture. First, to assess global ancestry, subjects from the 1000 Genomes Project were merged with the cleaned NHLP genotype data. The combined dataset was filtered for linkage disequilibrium (LD) in PLINK using the indep-pairwise method (Purcell et al., 2007). SNPs were then pruned from a 20-marker window until all markers had a pairwise R^2^ value below 0.1. The window was then slid two markers down the chromosome and the LD prune was repeated. In addition, known regions of long range LD were excluded (Price et al., 2008). This produced a final set of 239,414 SNPs for analysis. Finally, principal components analysis was performed using smartpca within the EIGENSOFT package (v6.1.4) with outlier removal enabled.(Price et al., 2006) Principal components were plotted in R to inspect separation and ensure the removal of regions of long range LD, which is a common confounder in analysis of admixed populations. A plot of PC1 versus PC2 is shown as Figure 2 to demonstrate admixture. Subjects that closely aligned with the East Asian cluster were excluded from downstream analysis (N = 4). Principal components were used to control for admixture in the primary GWAS.

**Figure 2.**
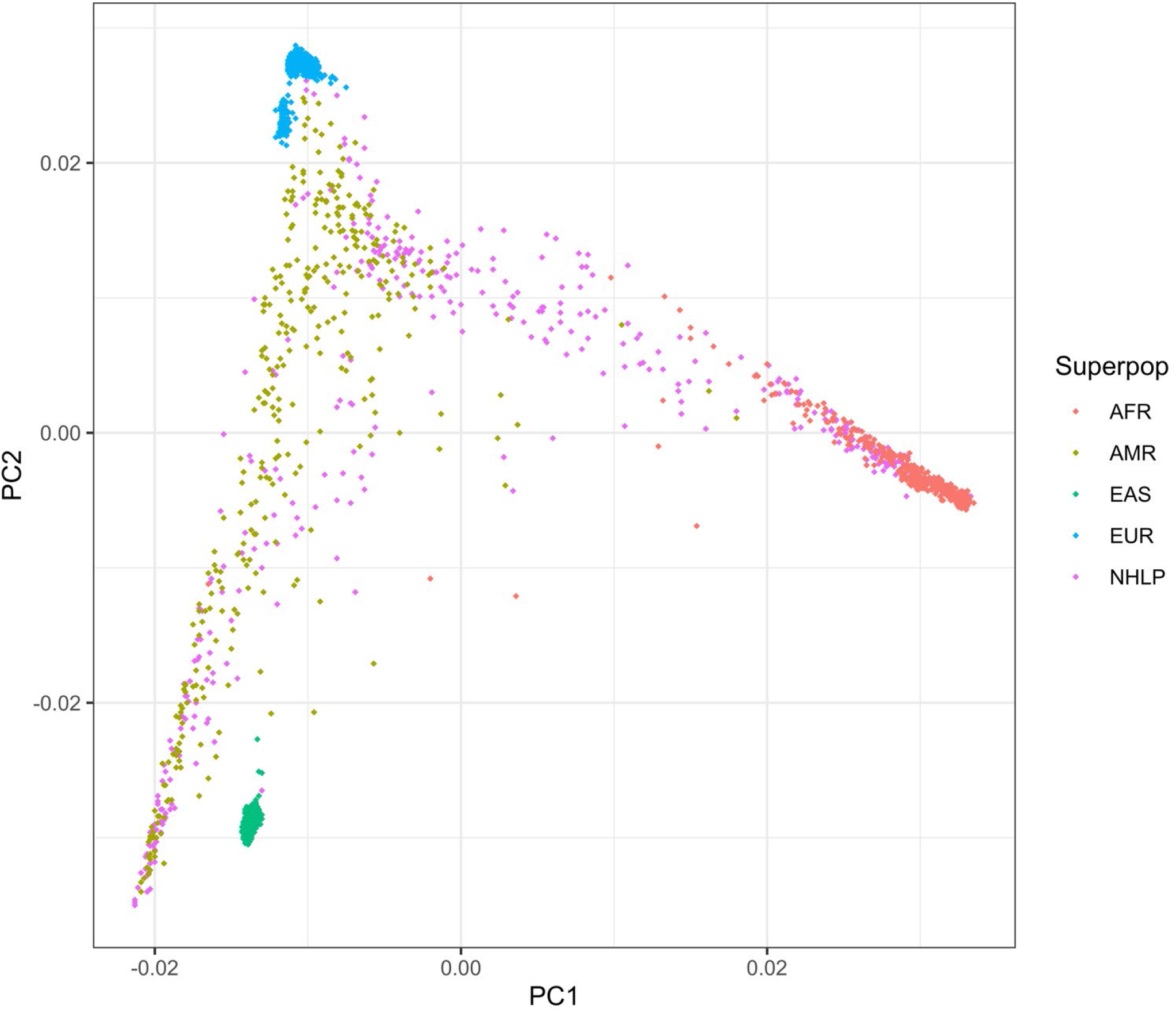
Plot of PC1 vs. PC2 for the NHLP merged with the 1000 Genomes Project. Plotting PC1 vs. PC2 shows population structure in the NHLP. PCA indicates that NHLP subjects are predominantly Hispanic or African American with small numbers of Europeans and Asians. AFR = African Superpopulation; AMR = Admixed American Superpopulation; EAS = East Asian Superpopulation; EUR = European Superpopulation.

### Primary GWAS Analysis

Logistic regressions were performed in PLINK v1.90-beta5.3 controlling for sex as encoded in the pedigree file, age at testing, ten ancestry principal components from EIGENSOFT, and the binary SES assignment.(Purcell et al., 2007) The ‘--adjusted’ flag was set to automatically calculate multiple testing comparisons and the lambda genomic inflation test statistic.

### *Post-hoc* NHLP Statistical Analysis

We performed three *post-hoc* tests to attempt to determine the likelihood of cryptic admixture potentially underlying the GWAS results. First, we used a 5×2 chi-square test to determine if there was an underlying difference in the proportion of minor and major alleles across self-report racial categories. Second, we used an ANOVA to compare the raw decoding composite scores to self-report racial categories to determine if scores were significantly different between categories. Finally, we performed Tukey’s honest significant differences test (Tukey’s HSD) to further check for pairwise differences between each self-report racial category and their performance on the decoding composite measure.

### *Tractor* Analysis

To attempt to identify the source ancestry that could account for the peak observed in the primary GWAS, we created ancestry specific VCF files from our admixed cohort using the software program *Tractor*.(Atkinson, 2020; Atkinson et al., 2020) First, autosomal genotype data were split by chromosome and phased using Eagle2 (v2.4.1).(Loh et al., 2016) For a reference panel, data from the 1000 Genomes Project were downloaded and converted to BCF using BCFtools (v1.5).(Li et al., 2009) Genetic maps were provided by the Eagle2 authors, REF/ALT swaps were allowed, and no imputation of missing alleles was performed. Following phasing, we performed local ancestry deconvolution using RFMIX.(Maples et al., 2013) For this, the full 1000 Genomes Project EUR (N = 503) and the continental AFR (N = 504) populations were used to capture European and African derived ancestral tracts.(The Genomes Project Consortium, 2015) To capture Native American ancestry present in Hispanic populations, we used the Peruvian from Lima population (PEL; N = 85) as they possess the highest overall fraction of this ancestry in the 1000 Genomes Project.(Martin et al., 2018) To account for the discrepancy between the number of subjects from the PEL population and the number of subjects from the other two populations, the terminal node size for the random forest trees in RFmix was set at 5. Next, we ran *Tractor* using the three-way admixed implementation scripts. This produced three separate ancestry-specific VCF files containing only ancestrally relevant segments of AFR, AMR, and EUR ancestry. We then performed separate logistic regression GWAS’s on these individual ancestry files for our dichotomized decoding composite, again using the covariates of age, sex, SES and ten PCs. This generated ancestry-specific effect size estimates and P-values for determining whether loci have differential effects in different ancestry groups.

### The Genes, Reading, and Dyslexia Study

Replication and follow-up analyses were performed in the Genes, Reading, and Dyslexia (GRaD) Study of 1,432 Hispanic American and African American subjects with microarray genotyping and psychoeducational testing. Genotyping, imputation, ancestry analysis, and covariates creation have been previously described (Truong, et al. 2019). For replication, we restricted our analysis to ages 7-9 years (N = 703), to match the developmental time window of the NHLP. Later analyses used 1,291 subjects across ages 7-16 years old. An analogous decoding latent variable was created using the same covariance modeling approach as described above for the NHLP sample. The following measures were used to form the latent variable in the GRaD sample: the WJ-III Word Attack, WJ-III Word Identification, TOWRE Sight Word Efficiency, TOWRE Nonword Decoding, and the Word Recognition Accuracy subtest of the Scholastic Reading Inventory (SRI). The GORT was not performed in the GRaD cohort; however, the Accuracy subtest of the SRI represents an equivalent construct of word decoding in context.

### Replication and Moderation Analysis in the GRaD Study

To replicate the primary GWAS results, we performed a logistic regression on the mean split decoding composite analogous to the composite created for the NHLP. GRaD imputed genotypes were filtered for candidate SNPs from the primary GWAS. Analysis was performed using PLINK v1.90-beta5.3 controlling for sex, age at testing, ten principal components from EIGENSOFT, and the binary SES assignment.(Purcell et al., 2007) The adjusted flag was set to calculate multiple testing statistics.

To investigate if age had a significant moderating influence on decoding performance we performed a moderation analysis in R v3.6.1.(R Core Team, 2016) SNPs were recoded under an additive model using PLINK. Using the untransformed decoding composite, we fit two linear regressions controlling for the same covariates as the logistic replication. In the first model, all variables were fit without any interaction terms. In the second model, an age-by-rs2599553 genotype interaction term was fit in addition to the main effect of each variable.

### Relative Risk Analysis

Subjects from the NHLP were identified as RD cases if they scored at least one standard deviation below developmental expectations on a minimum of two of three individually-administered and nationally-normed composite reading scores. The tests and composites used were as follows: WJ-III Basic Skills, WJ-III Broad Reading, TOWRE-II Index, and the GORT-5 Reading IQ. This criterion was applied to 323 subjects (79% of the full sample) for whom full assessment information was available at the start of Grade 2. Grade 2 was chosen as the target assessment point for establishing cases, since this represents the earliest date, in practice, that children at risk for reading failure can receive a full clinical assessment of reading skills.

### Growth Curves

Subjects from the NHLP were tested on a nationally normed reading assessment, the WJ-III, a maximum of nine times from the start of Grade 1 until the fall of Grade 5. Among the 412 children who completed at least one assessment point and had available genetic data, longitudinal data density was as follows: Grade 1 start, n = 383; Grade 1 end, n = 343; Grade 2 start, n = 368; Grade 2 end, n = 340; Grade 3 start, n = 380; Grade 3 end, n = 361; Grade 4 start, n = 359; Grade 4 end, n = 191; Grade 5 start, n = 174. Median number of assessments per child was seven, ranging from one to nine assessments. No differences in the proportion with the *GARRE1* risk allele were observed for longitudinal data density (*Χ^2^*(8) = 2.00, p = .90) or number of assessments (*Χ^2^*(8) = 5.09, p = .75) indicating that missing longitudinal data was not systematically related to the focal effect.

The following WJ-III subtests were used to formulate growth curves: Letter-Word Identification, measuring single-word identification; Word Attack, measuring orthographically-regular non-word decoding; Passage Comprehension, measuring reading of connected text for meaning via a cloze procedure; Reading Fluency, measuring both fluency and comprehension of connected text. Growth curves were then incorporated into models based on raw scores and standardized scores. Raw score models addressed absolute skill growth over time. Standard score models provided a picture representing how children changed relative to developmental expectations as characterized by the normative sample of the test. Standard scores on the WJ-III have a mean of 100, and a standard deviation of 15. In the standard score outcome analyses, a standard score of 100 was within developmental expectations; a standard score of 85 was one standard deviation below developmental expectations. A standard score below 85 is often used clinically to indicate significant problems acquiring reading skill, and as one of the criteria for diagnosing a reading-specific learning difficulty. A standard score of 90 is often used as an educational cut-off representing the low end of ‘average’ reading ability.

Growth curve models were formulated following best practices.(Hox et al., 2010; Snijders and Bosker, 2011) PROC MIXED in SAS/STAT software version 9.04 of the SAS System for Linux, was used to fit all multilevel growth models. After data screening and assessment of basic assumptions for distribution and outliers, the shape of individual growth trajectories was investigated empirically prior to analysis through visual inspection of each child’s trajectory from the fall of Grade 1 through the fall of Grade 5. Competitive approaches using different models of growth were then evaluated (e.g., linear versus higher-order versus growth to an asymptote, etc.). The most parsimonious and well-fitting growth model included an intercept centered at the Grade 1 start, with a linear growth component. Models of raw score test performance required an additional quadratic function to represent a general deceleration of growth rates over the observational period. Since the nine measurement timepoints have educational significance (i.e., beginning and end of each school year), but specific measurement dates varied per child, a hybrid model for time was implemented. Several models for time were considered against each other with a two-component model providing the best fit to the repeated measures elements in the model. In the first component, the nine fixed measurement occasions were modeled as random effects. In the second component, the number of days between measurements for each child was used to model the within-subject residual variance, using a spatial power covariance matrix.(Macchiavelli and Moser, 1997)

Sex, low versus average SES, and ten principal components to control for ancestry were included in the raw model as fixed-effect covariates for intercept, growth, and deceleration. No covariate predicted reading growth or deceleration, therefore all covariate by growth effects were pruned. The top SNP from the primary GWAS, rs2599553, was recoded for a dominance model and incorporated as follows: as a fixed-effect predictor of skill level differences across the study span; as a predictor of individual growth and change; and as a predictor of growth deceleration in the case of raw score models.

### Bioinformatic Analysis

To characterize the genes implicated by the GWAS results, we used publicly available expression quantitative trait loci (eQTL) and mRNA expression data from the Genotype-Tissue Expression Project (GTEx Project) and the Brainspan Project (Lonsdale et al., 2013; Miller et al., 2014). We only focused on expression data from brain tissues. The GTEx Project provides both eQTL data and mRNA expression from a limited number of brain regions in adult subjects. Brainspan data has fewer samples, but is designed to provide a more comprehensive look at gene expression in the brain over the human lifespan from early fetal development to middle age.

GTEx was interrogated through their website browser. Gene specific expression values and subject age were downloaded from the Brainspan portal for follow-up analysis. Expression was evaluated across developmental time in the Brainspan project in two broad life stages. Fetal samples (N = 11; ages 12 – 37 post-conception weeks) were divided into cerebellar and non-cerebellar. Differences in expression between the two region groupings were evaluated with the Wilcoxon’s Rank Sum test. This was repeated for cerebellar and non-cerebellar regions of the brain in the postnatal to adult samples (N = 21; ages 4 months – 40 yrs.).

The NIH-maintained LDlink server was used to investigate patterns of LD between the markers in the chromosome 19 peak. LD was calculated using the LDMatrix tool with the full African, European, and Admixed American 1000 Genomes Project superpopulations as reference.(Machiela and Chanock, 2015).

## Results

### Ancestry Analysis

Principal components analysis showed that the subjects in the NHLP were primarily of global majority race/ethnicities. When plotting PC1 vs. PC2 of the NHLP joined with subjects from the 1000 Genomes Project, we observed that our sample overlaps almost completely with subjects from the full AMR or full AFR superpopulations while a small group overlaps with subjects from the full EUR or full EAS superpopulations (Figure 2). This is consistent with a predominantly Hispanic and African American dataset, supported by self-reported racial category information.

### Primary GWAS

For the primary GWAS analysis in the NHLP, 407 subjects out of an initial 420 subjects with whole genome sequencing data were included. Four subjects were excluded for having self-report Asian ancestry. One subject was excluded for having a sibling in the dataset. Eight subjects lacked sufficient psycholoeducational testing data to generate the decoding composite score. No subjects were excluded for a lack of covariate data. Of the remaining 407 subjects, 179 subjects were assigned case (Z-score < 0) status and 228 were assigned control (Z-score ≥ 0) status (Figure 1).

Logistic regression based GWAS of the latent variable derived case/control status identified a cluster of chromosome 19 SNPs centered on and around the gene called *GARRE1* (Figure 3A). Of the SNPs in this cluster, a single SNP (rs2599553), demonstrated a P-value that exceeded the conventional genome-wide significance threshold of 5 x 10^−8^ (Figure 3A). All 31 genome-wide significant or suggestive SNPs from the primary GWAS are included in Table 3. The lowest observed P-value was 3.13 x 10^−8^ with an odds ratio of 3.381 for rs2599553 (Table 3). Manual inspection of the Quantile-Quantile (Q-Q) plot (Figure 3B), and the calculated lambda test statistic for inflation of 1.04149, both support the assertion that the primary GWAS was well-controlled for confounding due to admixture.

**Figure 3.**
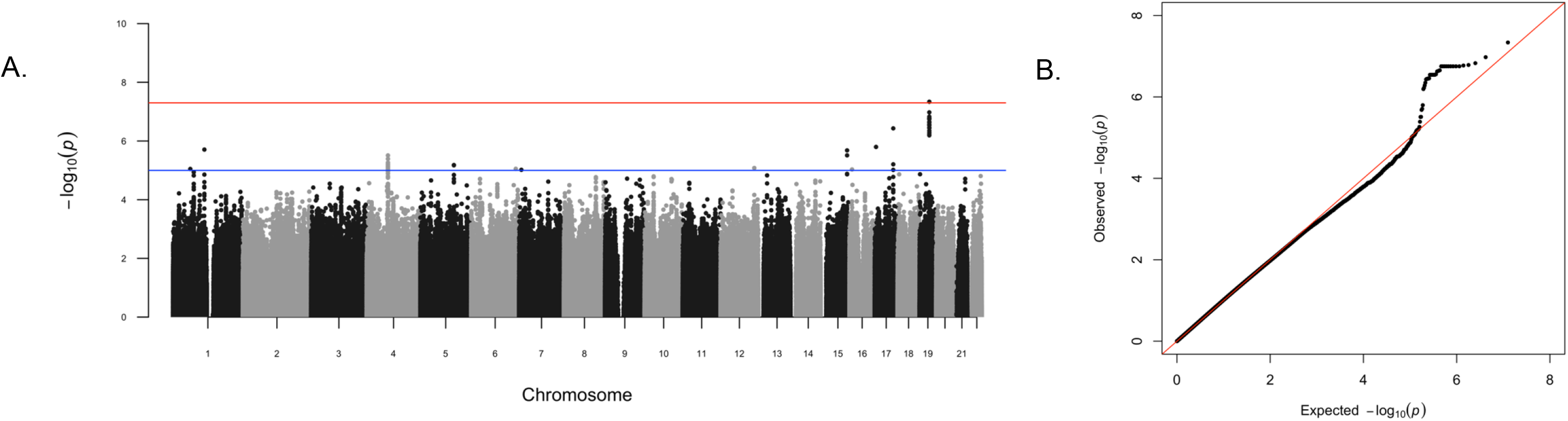
Panel A: Manhattan plot of primary GWAS in the NHLP. The red line indicates the genome wide significance threshold of 5×10^−8^. The blue line indicates a suggestive threshold of 1×10^5^. P-values are negative log transformed. Panel B: Q-Q Plot corresponding to GWAS results in Panel A.

**Table 3.** Primary GWAS results for chromosome 19 sorted by base pair position in the NHLP. Significant or suggestive or significant SNPs are reported. BP is basepair in hg19 coordinates. OR is odds ratio from PLINK. Top SNPs rs2599553 is highlighted in yellow.

| rsID | BP | Minor Allele | OR | P-Value |
| --- | --- | --- | --- | --- |
| rs2115487 | 34742162 | A | 3.184 | 1.52E-07 |
| rs11084762 | 34750567 | G | 2.988 | 1.88E-07 |
| rs7359931 | 34754797 | G | 3.247 | 9.22E-08 |
| rs4805079 | 34757224 | A | 3.247 | 9.22E-08 |
| rs10426700 | 34764563 | T | 3.247 | 9.22E-08 |
| rs10407101 | 34765113 | T | 3.247 | 9.22E-08 |
| rs10407640 | 34766588 | G | 3.247 | 9.22E-08 |
| rs12975032 | 34769256 | C | 3.217 | 1.19E-07 |
| rs11671239 | 34775721 | A | 3.247 | 9.22E-08 |
| rs328412 | 34800749 | A | 3.247 | 9.22E-08 |
| rs328414 | 34802393 | A | 3.278 | 7.49E-08 |
| rs1664905 | 34805294 | T | 3.247 | 9.22E-08 |
| rs921476 | 34807200 | A | 3.247 | 9.22E-08 |
| rs2599553 | 34814089 | A | 3.381 | 3.13E-08 |
| rs1669263 | 34816031 | C | 3.316 | 5.47E-08 |
| rs2965269 | 34819331 | A | 3.257 | 1.27E-07 |
| rs189030 | 34820159 | C | 3.165 | 1.48E-07 |
| rs1618249 | 34826702 | T | 3.214 | 1.20E-07 |
| rs328402 | 34829261 | C | 3.165 | 1.48E-07 |
| rs7254168 | 34834364 | A | 3.272 | 9.29E-08 |
| rs328406 | 34838998 | T | 3.165 | 1.48E-07 |
| rs328405 | 34840634 | A | 3.212 | 1.23E-07 |
| rs62122220 | 34841826 | A | 3.165 | 1.48E-07 |
| rs397072 | 34847014 | A | 3.165 | 1.48E-07 |
| rs422732 | 34848450 | C | 3.165 | 1.48E-07 |
| rs416602 | 34851509 | C | 3.167 | 1.70E-07 |
| rs8191356 | 34855095 | A | 3.167 | 1.70E-07 |
| rs7260568 | 34892284 | A | 3.203 | 1.58E-07 |
| rs35024640 | 34940263 | T | 3.186 | 1.34E-07 |
| rs16969326 | 34940644 | T | 3.135 | 2.00E-07 |
| rs60857340 | 34943280 | T | 3.186 | 1.34E-07 |

### Post hoc Analysis of rs2599553 and the Decoding Composite

To test for possible confounding due to variability in allele frequencies between self-reported racial categories in the NHLP, we performed a post-hoc 5×2 chi-square test (Table 4). This showed a test statistic of 1.0391 with a corresponding P-value of 0.90381, which does not support a significant deviation in allele counts between self-reported racial groupings (χ^2^(4, 407) = 1.0391, P = n.s.). To test for possible confounding due to variability in the phenotype between self-report racial categories, we performed a one-way ANOVA (Table 5) and Tukey’s HSD test. Box-plots of raw decoding composite scores were plotted by self-report racial category (Figure 4). The ANOVA showed an F-test statistic of 2.07 and a corresponding P-value of 0.084, indicating no statistically significant differences in decoding score by self-report racial groupings (F(4,402) = 2.07, *p* = 0.084; Table 5). Tukey’s HSD tests for differences in means between each pair of self-report racial categories, and is generally applied as a follow-up to ANOVA if results are significant. Despite no significant association between self-ID racial category and performance on the decoding composite, we performed Tukey’s HSD. No significant effects were observed for any pair of self-reported racial categories. Although confounding due to variability in allele frequency or phenotype is the most common cause of P-value inflation in GWAS’s, these studies show that it is not a significant confounder in the present study.

**Figure 4.**
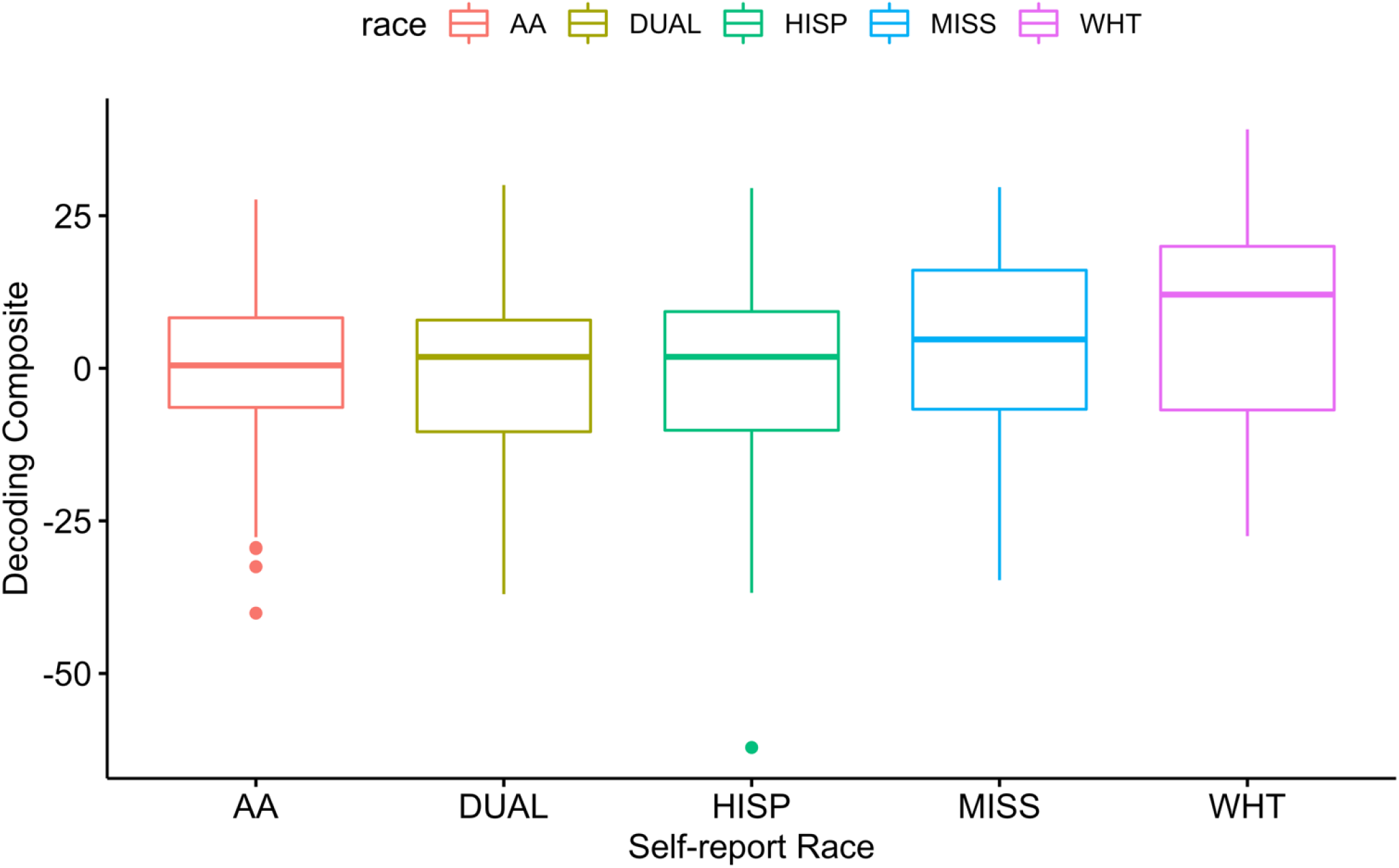
Raw decoding composite scores for each self-report racial grouping (N=415) in the NHLP. DUAL indicates more than one category selected, MISS indicates no data at this question.

**Table 4.** Chi-square test for difference of minor/major allele counts across self-report identities in the NHLP.

| rs2599553 | AA (N = 95) | HISP (N = 107) | WHT (N = 19) | DUAL (N= 99) | NA (N = 96) |  |
| --- | --- | --- | --- | --- | --- | --- |
| Minor | 30 | 41 | 7 | 38 | 36 | Test Statistic = 1.0391<br>P = 0.90381 |
| Major | 160 | 173 | 31 | 160 | 156 |  |
| MAF | 0.158 | 0.192 | 0.184 | 0.192 | 0.1875 |  |

**Table 5.** One-way ANOVA results for differences in mean across self-report racial groupings in the NHLP.

|  | DF | Sum Sq | Mean Sq | F-Test Value | P-Value |
| --- | --- | --- | --- | --- | --- |
| Self-Report Race | 4 | 2014 | 503.5 | 2.07 | 0.084 |
| Residuals | 402 | 97809 | 243.5 |  |  |

### *Tractor* Analysis

*Tractor* was used to partition the phased, joint-called NHLP genotype files into three separate VCF files corresponding to the African, European, and Admixed American ancestry-specific haplotype tracts. Individual GWAS’s for each ancestry of the N=415 subjects were performed using the covariates described above. Individual ancestry GWAS indicated that the segment on chromosome 19 that had the strongest association with decoding performance in the primary GWAS, had the strongest signal in AFR ancestry (rs2599553; P = 0.000457, OR=3.339; Figure 5A). For comparison, AMR ancestry also showed an association with P = 0.005229, and OR=2.557 (Figure 5B) while EUR ancestry only showed a nonsignificant P-value of 0.4398, and OR = 1.217 (Figure 5C). These differences in P-values suggest that the signal for this genetic variant in the primary GWAS mostly came from the non-European ancestry in the admixed sample and highlight the utility of accounting for local ancestry in admixed cohorts.

**Figure 5:**
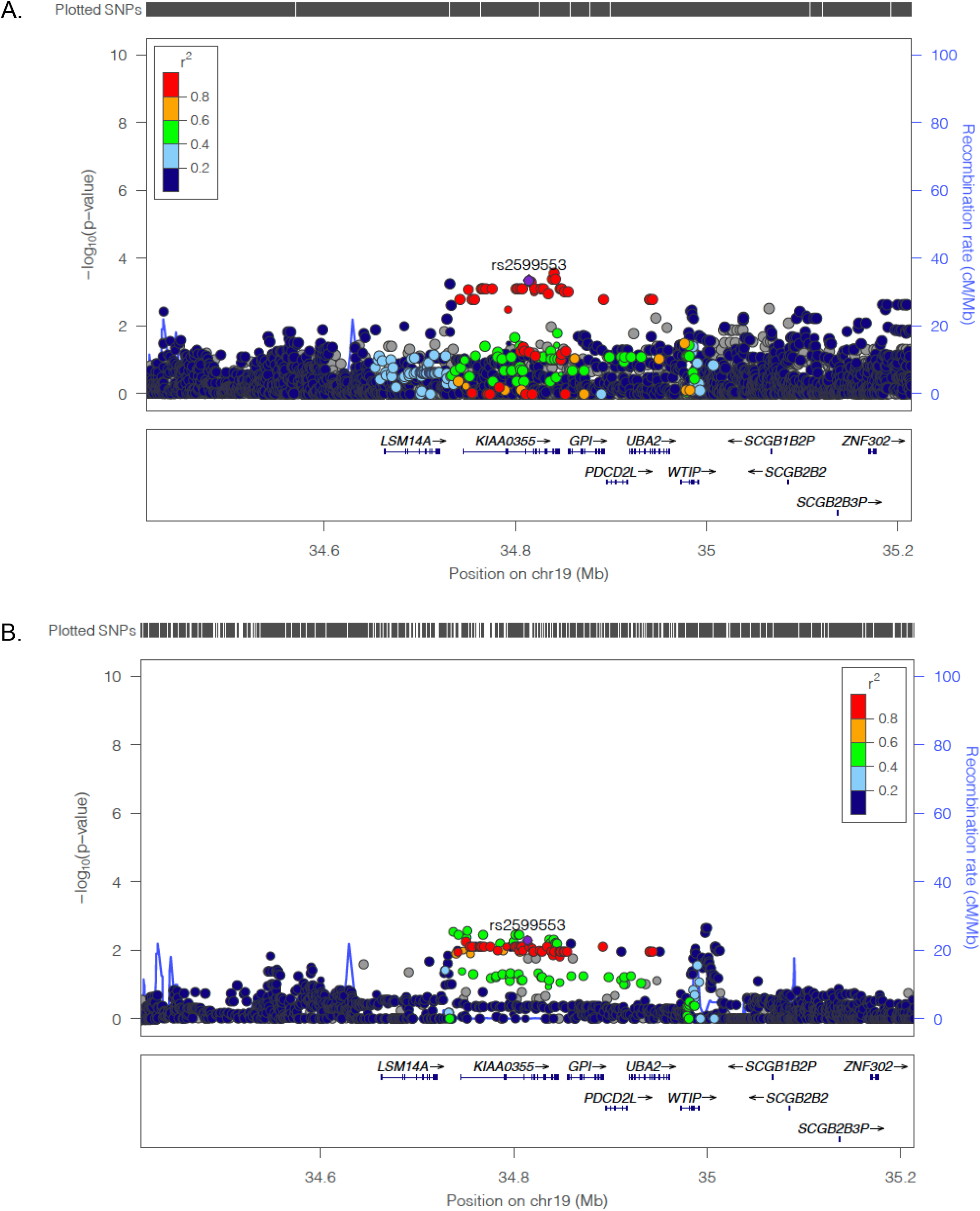
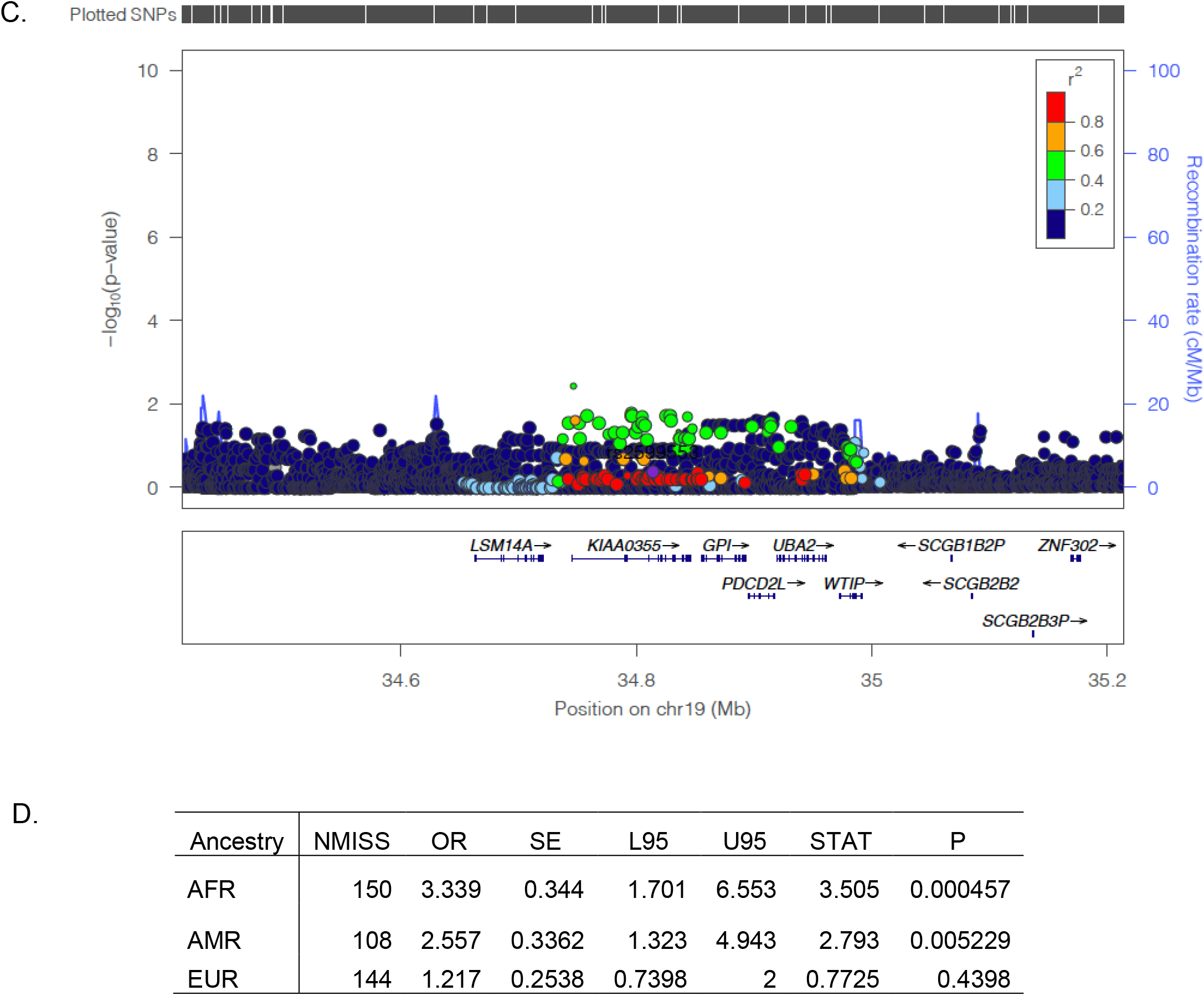
Ancestry-specific GWAS results for the chr19 locus highlighting lead SNP rs2599553 (purple diamond), in the NHLP. The color of SNPs corresponds to the level of correlation with rs2599553 using the 1000 Genomes Project EUR LD map. SNP location and density is visualized at the top of the image. Genes in the area are visualized in the bottom track. Panel A are the results in the AFR population, Panel B are the results for AMR, Panel C are the results for EUR, Panel D are the summary statistics from the individual GWAS and the meta-analysis.

### Bioinformatic Analysis

The 31 SNPs comprising the chromosome 19 peak span 250 Kbp and four non-overlapping genes: *GARRE1*, *GPI*, *PDCDL2*, and *UBA2*. All 31 SNPs are non-coding, and 22 overlap *GARRE1*. LD analysis showed that all 31 SNPs had R^2^ values above 0.95, indicating a single locus, and could not be used to differentiate between the four genes. Thirty SNPs were observed in the GTEx eQTL dataset, and all were eQTLs for *GARRE1* expression in the cerebellum. The lack of eQTL evidence for any other of the genes in the chromosome 19 peak strongly implicates *GARRE1* as a candidate gene for decoding performance. Bulk mRNA sequencing from the GTEx project showed peak brain expression in the cerebellum with a median TPM value for *GARRE1* of 23.71 in cerebellar hemisphere and 21.86 in cerebellum (cerebellar hemisphere and cerebellum are treated as replicates sampled at two separate times by two separate teams), supporting the eQTL observations.

Brainspan data showed that in human fetal samples between 12 to 37 weeks post-conception (Figure 6A), *GARRE1* expression in the cortex and cerebellum are equivalent (Wilcoxon’s Rank Sum test p = 0.12; N = 11 subjects). However, beginning at 4 months postnatal age and extending well into adulthood there is significantly higher expression of *GARRE1* in the cerebellum relative to all other brain tissues (Wilcoxon’s Rank Sum test p = 6.88 x 10^−11^; N = 21 subjects)(Figure 6B).

**Figure 6.**
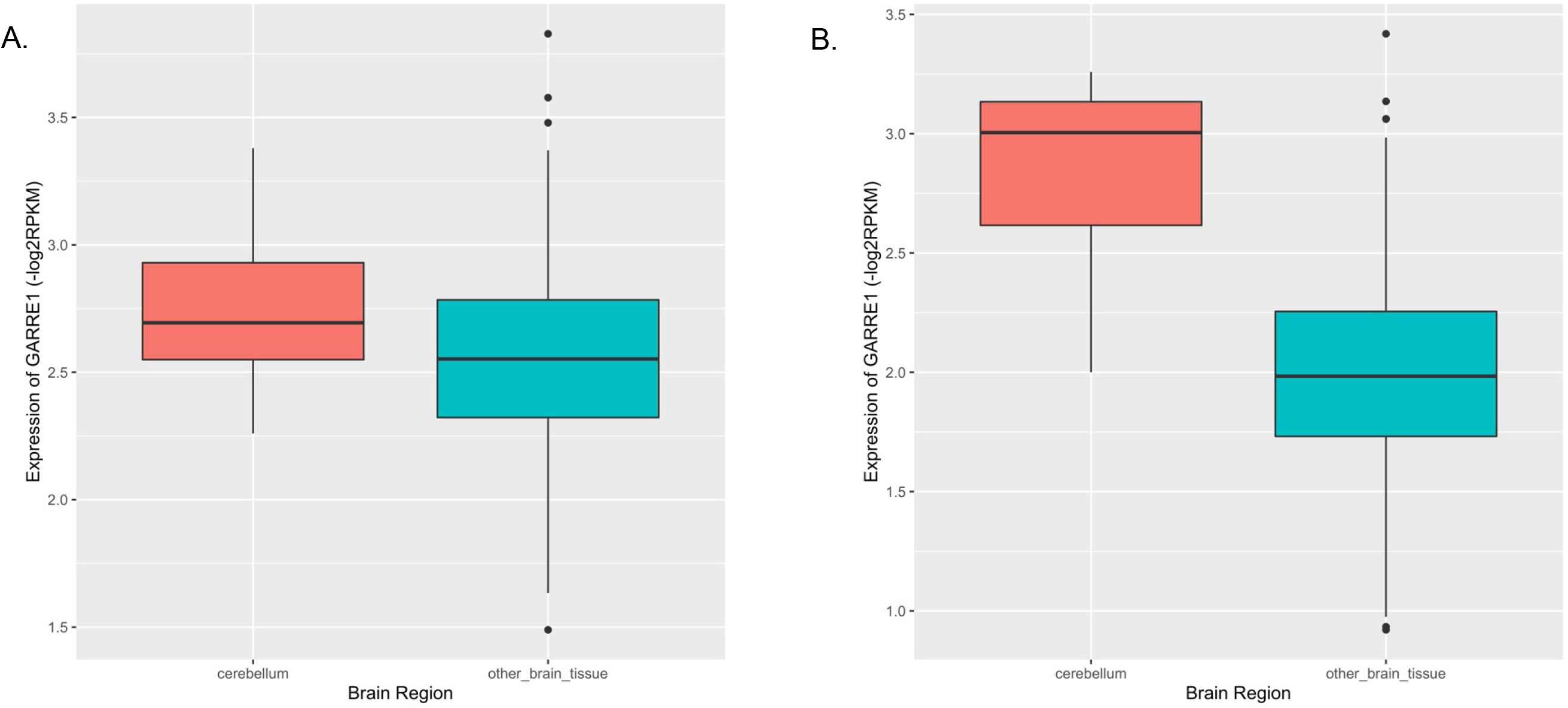
Brainspan expression of *GARRE1*. Panel A shows expression of *GARRE1* for all fetal samples (12-37pcw, N = 11). Panel B shows expression of *GARRE1* for post-natal samples (4 mo.- 40 yrs., N = 21). Expression data suggest relatively constant expression of *GARRE1* in the cerebellum prenatally and postnatally, while expression in other regions of the brain drops postnatally.

Data from the gnomAD project suggested that *GARRE1* is intolerant to loss of function (pLOF) mutations with a pLI score of 0.97 and a ratio of observed to expected pLOF mutations of 0.17. The expectation under a neutral model is that 46.4 pLOF mutations would be observed in a dataset the size of gnomAD, however, only 8 were observed for *GARRE1*. In contrast, fewer missense mutations were observed (507) compared to the 622 that were expected. The number of synonymous mutations fell within the expected ratio, with 272 observed compared to an expected 254.6. Together these data indicate that *GARRE1* is performing an important function in humans that requires two intact genes for successful reproduction.

### Replication in GRaD

For replication, we chose the GRaD Study because subjects were assessed with a robust battery that included single-word decoding skills, they were previously genotyped with a large number of SNPs, and because the GRaD sample has a broad representation of Hispanic-American and African-American children from different regions of the US. Using the same set of covariates from the primary GWAS in the NHLP and an analogous mean split decoding composite, we replicated our lead SNP (N = 632; Table 6). The best performing SNP in the NHLP, rs2599553, had a P-value of 0.015 in GRaD; the best results in GRaD were from rs2965269, P-value = 0.012. The pairwise R^2^ values were above 0.95 for all 31 SNPs, suggesting that there is only a single effective test, avoiding the need for a multiple testing correction. Interestingly, we were only able to replicate when we age-matched GRaD subjects to NHLP (restricting the GRaD to 7-10 year old subjects), before mean splitting the composite, suggesting a possible gene-by-age effect.

**Table 6.**
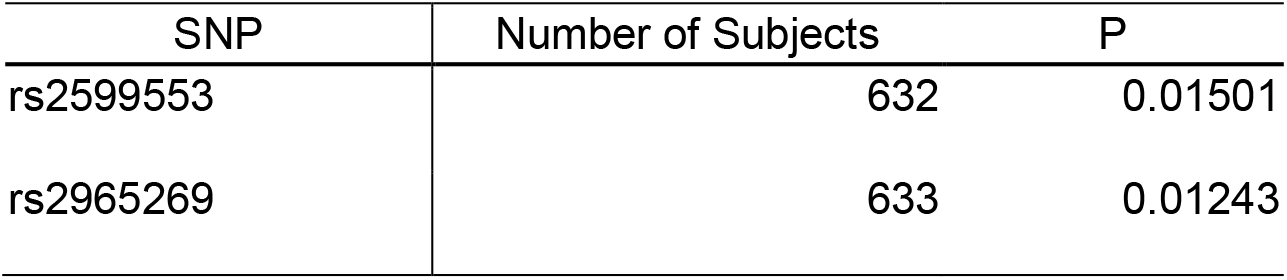
GRaD Candidate SNP replication. Columns are SNP ID, number of subjects included in the model and P-value from logistic regression.

| SNP | Number of Subjects | P |
| --- | --- | --- |
| rs2599553 | 632 | 0.01501 |
| rs2965269 | 633 | 0.01243 |

### Moderation Analysis

To investigate a potential gene-by-age effect, we performed a SNPxAGE moderation analysis for the lead SNP (rs2599553) from the primary analysis in NHLP. Using the quantitative, normally distributed, decoding composite for the full GRaD cohort (N = 1,291), we performed a regression with and without a rs2599553xAge interaction term. Both models included age, sex, a binary SES variable, and ten PCs to control for admixture. The main effect of rs2599553 genotype was significant only when the interaction term was included in the model (P < 0.05); the P-value for rs2599553xAge was also less than 0.05 (Table 7). These analyses indicate that age had a significant moderating influence on the effect of rs2599553 on decoding performance. Stratifying age by quantile, increasing numbers of the minor allele was associated with a positive direction of effect on decoding skill at ages 7 through 8 years, and a negative direction of effect at ages 13 through 16 years. The central 50% of the age/performance distribution curve, ages 9 through 12, was relatively flat and showed no relationship between presence of minor alleles and decoding skill (Figure 7).

**Figure 7.**
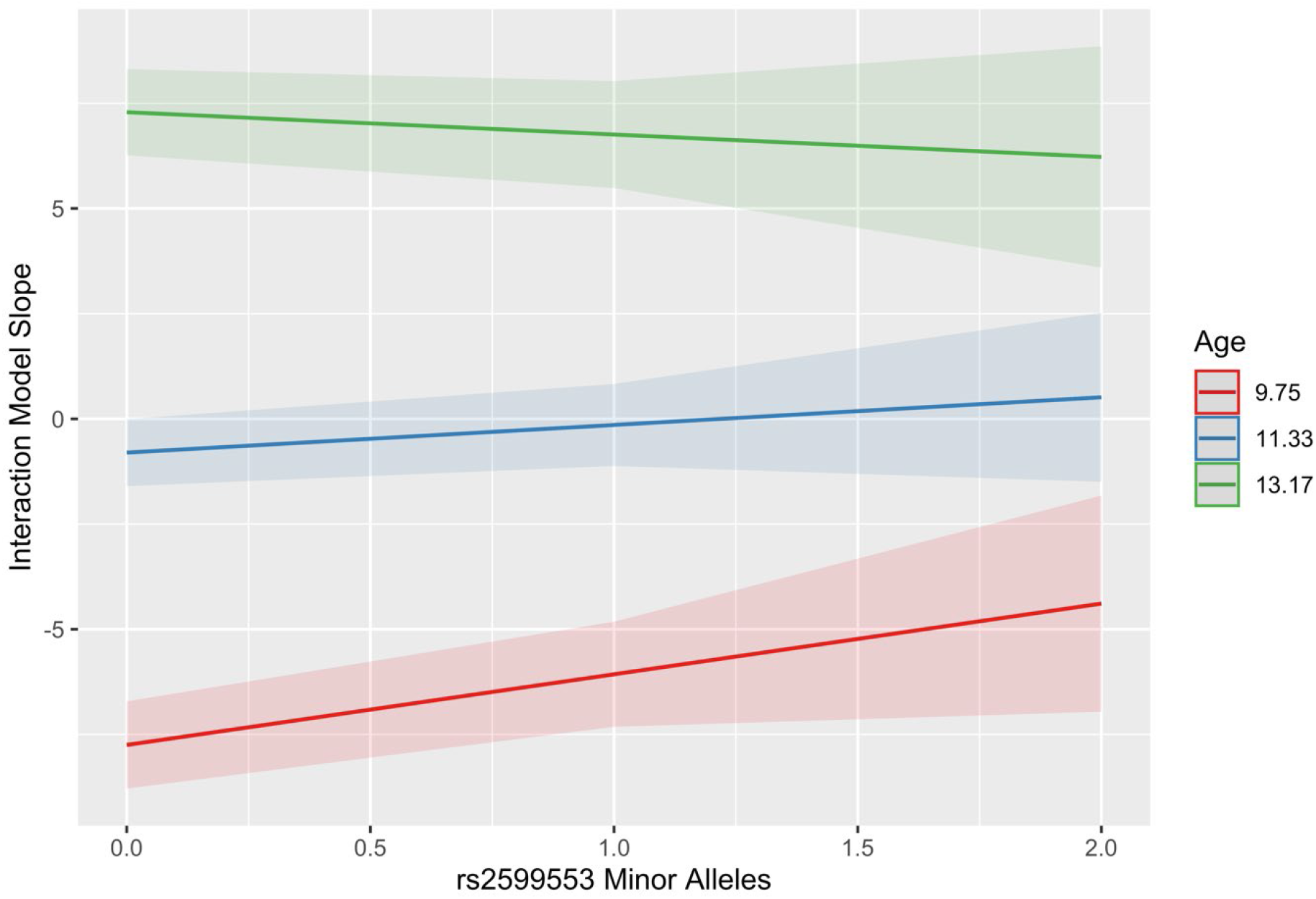
Marginal slopes plot for minor alleles of rs2599553 and decoding performance. The red line represents the slope of the regression for the bottom 25% of subjects in the GRaD by age, the blue line represents the slope of the regression for the center 50% of subjects in the GRaD by age, and the green line represents the slope of the regression for the top 25% of subjects by age.

**Table 7.** GRaD Moderation analysis results. In the model summarized in the left column, there is no interaction term. Age is not a significant predictor of decoding performance. In the model summarized in the right column, the SNP by decoding relationship is moderated by age. Both the SNP and the moderation term are significant, suggesting age moderates the relationship between rs2599553 and decoding.

|  | *Dependent variable:* |  |
| --- | --- | --- |
|  | GRaD Decoding Composite |  |
|  | No Interaction | Age/SNP Interaction |
| Constant | -47.464*** (-51.394, -43.534) | -50.308*** (-54.952, -45.665) |
| rs2599553_A | 0.603 (-0.578, 1.783) | 7.988* (1.434, 14.541) |
| age | 4.148*** (3.832, 4.463) | 4.398*** (4.015, 4.781) |
| SEX | 2.164** (0.850, 3.477) | 2.207** (0.895, 3.519) |
| SES_CC | -2.617*** (-4.020, -1.213) | -2.694*** (-4.097, -1.291) |
| EV1 | 37.469** (13.421, 61.517) | 37.505** (13.495, 61.516) |
| EV2 | 61.280*** (36.189, 86.370) | 61.142*** (36.091, 86.193) |
| EV3 | -10.597 (-35.025, 13.831) | -11.148 (-35.542, 13.246) |
| EV4 | 5.522 (-18.165, 29.209) | 5.088 (-18.564, 28.740) |
| EV5 | 4.959 (-18.652, 28.570) | 4.100 (-19.486, 27.685) |
| EV6 | -3.449 (-27.094, 20.196) | -3.518 (-27.125, 20.090) |
| EV7 | -3.085 (-26.767, 20.597) | -2.285 (-25.940, 21.370) |
| EV8 | 21.818 (-1.737, 45.372) | 23.119 (-0.425, 46.664) |
| EV9 | -11.444 (-35.211, 12.324) | -11.489 (-35.219, 12.241) |
| EV10 | 19.076 (-4.507, 42.658) | 18.708 (-4.839, 42.256) |
| rs2599553_A:age |  | -0.647* (-1.212, -0.082) |
| Observations | 1,291 | 1,291 |
| R<sup>2</sup> | 0.402 | 0.405 |
| Adjusted R<sup>2</sup> | 0.396 | 0.398 |
| Residual Std. Error | 11.962 (df = 1276) | 11.943 (df = 1275) |
| F Statistic | 61.315*** (df = 14; 1276) | 57.745*** (df = 15; 1275) |

*Note:*
\* p<0.05; \*\* p<0.01; \*\*\* p<0.001

### Relative Risk

The top SNP from the primary GWAS, rs2599553, was coded according to a dominance model and used to calculate the relative risk of case status. Of the 323 subjects in this analysis, 101 were coded as having risk due to minor alleles of rs2599553, and of those, 39.6% (n = 40) were RD cases. 222 subjects were coded as having no allele risk, and of those, 21.2% (n = 47) were RD cases. Taken together, having the minor allele of rs2599553 conferred a 2.11 relative risk for meeting RD criteria at the start of Grade 2, assuming a conservative prevalence of 11% for reading disability in the general population.(Fletcher et al., 2007). Expressed differently, the top SNP from the primary GWAS conferred a 111% elevated risk of meeting the criteria for RD in Grade 2.

### Growth Curve Analysis

Across all four raw score growth models, substantial child-to-child variability was observed in the random effects for intercept, growth rates, and deceleration, indicating that growth over time was not influenced by the timing of when a child was enrolled, in Grade 1 or 2 (*data not shown)*. The general pattern of growth in reading skill was increasing raw score ability that decelerated over the study period (Figure 8a). Children with the *GARRE1* risk allele had consistently suppressed reading skill, compared to those without. This effect was replicated across all four reading measures and across all observation points from Grade 1 to Grade 5. The estimated difference in mean raw reading scores between risk and no-risk groups is consistently significant across all ages and reading measures (Table 8).

**Figure 8a.**
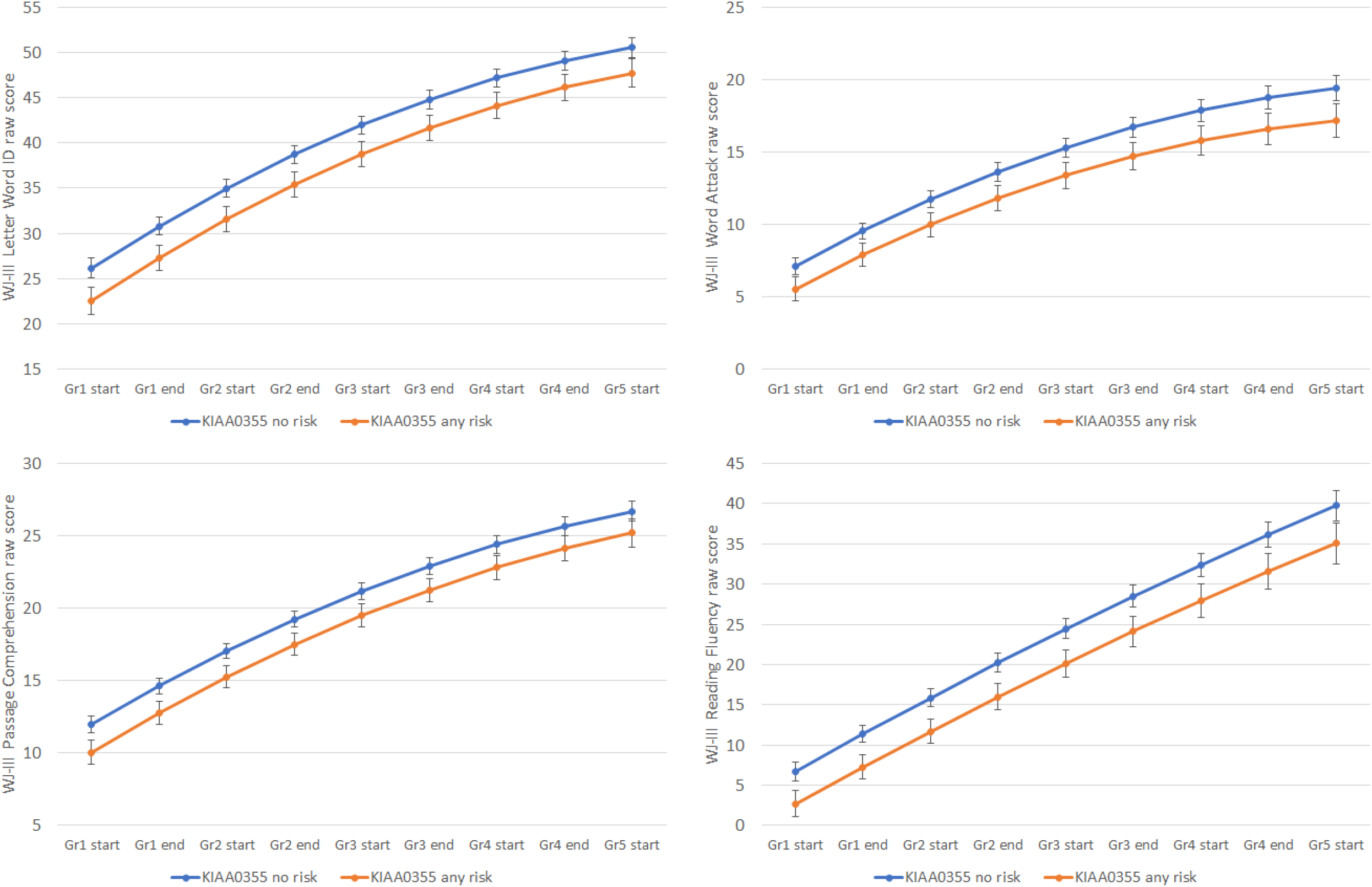
Growth curves depicting model-estimated raw scores on four standardized reading assessments from the Woodcock-Johnson III, and covering the developmental period from the start of Grade 1 until the start of Grade 5. Within each graph, growth curves are presented separately for the two GARRE1 risk groups.

**Table 8.** Woodcock-Johnson III estimated raw score mean differences between GARRE1 risk and no-risk groups in the NHLP.

| Subscale | Timepoint | Estimate | SE | P-value | Upper 95% CI | Lower 95% CI |
| --- | --- | --- | --- | --- | --- | --- |
| Letter Word ID | Grade 1 Start | 3.6 | 0.93 | 0.0001 | 1.8 | 5.5 |
|  | Grade 1 End | 3.5 | 0.88 | <.0001 | 1.8 | 5.3 |
|  | Grade 2 Start | 3.4 | 0.85 | <.0001 | 1.8 | 5.1 |
|  | Grade 2 End | 3.3 | 0.84 | <.0001 | 1.7 | 5.0 |
|  | Grade 3 Start | 3.2 | 0.84 | 0.0001 | 1.6 | 4.9 |
|  | Grade 3 End | 3.1 | 0.85 | 0.0003 | 1.5 | 4.8 |
|  | Grade 4 Start | 3.0 | 0.87 | 0.0006 | 1.3 | 4.7 |
|  | Grade 4 End | 2.9 | 0.91 | 0.0015 | 1.1 | 4.7 |
|  | Grade 5 Start | 2.8 | 0.96 | 0.0036 | 0.93 | 4.7 |
| Word Attack | Grade 1 Start | 1.6 | 0.51 | 0.0021 | 0.57 | 2.6 |
|  | Grade 1 End | 1.7 | 0.49 | 0.0009 | 0.69 | 2.6 |
|  | Grade 2 Start | 1.7 | 0.49 | 0.0005 | 0.77 | 2.7 |
|  | Grade 2 End | 1.8 | 0.51 | 0.0004 | 0.83 | 2.8 |
|  | Grade 3 Start | 1.9 | 0.53 | 0.0004 | 0.86 | 3.0 |
|  | Grade 3 End | 2.0 | 0.57 | 0.0006 | 0.86 | 3.1 |
|  | Grade 4 Start | 2.1 | 0.62 | 0.0009 | 0.85 | 3.3 |
|  | Grade 4 End | 2.2 | 0.68 | 0.0016 | 0.83 | 3.5 |
|  | Grade 5 Start | 2.2 | 0.74 | 0.0026 | 0.79 | 3.7 |
| Passage Comprehension | Grade 1 Start | 1.9 | 0.51 | 0.0002 | 0.91 | 2.9 |
|  | Grade 1 End | 1.9 | 0.48 | 0.0001 | 0.91 | 2.8 |
|  | Grade 2 Start | 1.8 | 0.47 | 0.0001 | 0.89 | 2.7 |
|  | Grade 2 End | 1.8 | 0.46 | 0.0002 | 0.84 | 2.7 |
|  | Grade 3 Start | 1.7 | 0.47 | 0.0003 | 0.77 | 2.6 |
|  | Grade 3 End | 1.6 | 0.49 | 0.0008 | 0.68 | 2.6 |
|  | Grade 4 Start | 1.6 | 0.52 | 0.0023 | 0.57 | 2.6 |
|  | Grade 4 End | 1.5 | 0.55 | 0.006 | 0.45 | 2.6 |
|  | Grade 5 Start | 1.5 | 0.60 | 0.014 | 0.31 | 2.7 |
| Reading Fluency | Grade 1 Start | 4.0 | 0.98 | <.0001 | 2.1 | 5.9 |
|  | Grade 1 End | 4.1 | 0.95 | <.0001 | 2.2 | 6.0 |
|  | Grade 2 Start | 4.2 | 0.96 | <.0001 | 2.3 | 6.0 |
|  | Grade 2 End | 4.2 | 1.00 | <.0001 | 2.3 | 6.2 |
|  | Grade 3 Start | 4.3 | 1.07 | <.0001 | 2.2 | 6.4 |
|  | Grade 3 End | 4.4 | 1.17 | 0.0002 | 2.1 | 6.7 |
|  | Grade 4 Start | 4.5 | 1.28 | 0.0005 | 2.0 | 7.0 |
|  | Grade 4 End | 4.6 | 1.42 | 0.0014 | 1.8 | 7.4 |
|  | Grade 5 Start | 4.7 | 1.56 | 0.0031 | 1.6 | 7.7 |

Standard score growth models depict a different perspective, placing each child’s score in relation to developmental expectations. In the standard score growth models, risk group was a significant predictor of ability for all dimensions of reading: Letter Word Identification, Word Attack, Passage Comprehension, and Reading Fluency. These differences in risk group reading performance were consistently significant at all time points (Table 9). Since the standard deviation of the standard scores for these measures is 15, these data show that the developmental risk conferred by *GARRE1* ranges from one-third (Word Attack outcome) to over one-half of a standard deviation (Reading Fluency outcome) below age expectations for reading performance.

**Table 9.** Woodcock-Johnson III estimated standard score mean differences between GARRE1 risk and no-risk groups in the NHLP.

| Subscale | Timepoint | Estimate | SE | P-value | Upper 95% CI | Lower 95% CI |
| --- | --- | --- | --- | --- | --- | --- |
| Letter Word ID | Grade 1 Start | 5.7 | 1.7 | 0.0007 | 2.4 | 9.0 |
|  | Grade 1 End | 5.6 | 1.6 | 0.0005 | 2.4 | 8.7 |
|  | Grade 2 Start | 5.4 | 1.5 | 0.0004 | 2.4 | 8.4 |
|  | Grade 2 End | 5.3 | 1.5 | 0.0004 | 2.4 | 8.2 |
|  | Grade 3 Start | 5.2 | 1.5 | 0.0005 | 2.3 | 8.1 |
|  | Grade 3 End | 5.0 | 1.5 | 0.0007 | 2.1 | 7.9 |
|  | Grade 4 Start | 4.9 | 1.5 | 0.0013 | 1.9 | 7.9 |
|  | Grade 4 End | 4.8 | 1.6 | 0.0024 | 1.7 | 7.8 |
|  | Grade 5 Start | 4.6 | 1.6 | 0.0047 | 1.4 | 7.8 |
| Word Attack | Grade 1 Start | 3.1 | 1.3 | 0.022 | 0.45 | 5.7 |
|  | Grade 1 End | 3.2 | 1.3 | 0.012 | 0.71 | 5.6 |
|  | Grade 2 Start | 3.3 | 1.2 | 0.0062 | 0.94 | 5.6 |
|  | Grade 2 End | 3.4 | 1.2 | 0.0035 | 1.1 | 5.7 |
|  | Grade 3 Start | 3.5 | 1.1 | 0.0022 | 1.3 | 5.8 |
|  | Grade 3 End | 3.6 | 1.2 | 0.0018 | 1.4 | 5.9 |
|  | Grade 4 Start | 3.8 | 1.2 | 0.0017 | 1.4 | 6.1 |
|  | Grade 4 End | 3.9 | 1.2 | 0.0019 | 1.4 | 6.3 |
|  | Grade 5 Start | 4.0 | 1.3 | 0.0025 | 1.4 | 6.6 |
| Passage Comprehension | Grade 1 Start | 5.2 | 1.5 | 0.0005 | 2.3 | 8.0 |
|  | Grade 1 End | 4.9 | 1.4 | 0.0004 | 2.2 | 7.6 |
|  | Grade 2 Start | 4.6 | 1.3 | 0.0005 | 2.1 | 7.2 |
|  | Grade 2 End | 4.4 | 1.3 | 0.0007 | 1.9 | 6.9 |
|  | Grade 3 Start | 4.1 | 1.3 | 0.0013 | 1.6 | 6.6 |
|  | Grade 3 End | 3.8 | 1.3 | 0.0032 | 1.3 | 6.4 |
|  | Grade 4 Start | 3.6 | 1.3 | 0.0085 | 0.91 | 6.2 |
|  | Grade 4 End | 3.3 | 1.4 | 0.022 | 0.49 | 6.1 |
|  | Grade 5 Start | 3.0 | 1.5 | 0.049 | 0.012 | 6.0 |
| Reading Fluency | Grade 1 Start | 8.7 | 2.0 | <.0001 | 4.8 | 12.6 |
|  | Grade 1 End | 8.3 | 1.8 | <.0001 | 4.7 | 11.8 |
|  | Grade 2 Start | 7.8 | 1.6 | <.0001 | 4.6 | 11.0 |
|  | Grade 2 End | 7.4 | 1.5 | <.0001 | 4.4 | 10.4 |
|  | Grade 3 Start | 7.0 | 1.5 | <.0001 | 4.1 | 9.9 |
|  | Grade 3 End | 6.5 | 1.5 | <.0001 | 3.6 | 9.4 |
|  | Grade 4 Start | 6.1 | 1.5 | <.0001 | 3.1 | 9.1 |
|  | Grade 4 End | 5.7 | 1.6 | 0.0006 | 2.4 | 8.9 |
|  | Grade 5 Start | 5.2 | 1.8 | 0.0036 | 1.7 | 8.8 |

**Table 10.** Random Effects Covariance Parameter Estimates for Woodcock-Johnson III Raw Scores.

|  | **Letter Word ID** |  |  | **Word Attack** |  |  | **Passage Comprehension** |  |  | **Reading Fluency** |  |  |
| --- | --- | --- | --- | --- | --- | --- | --- | --- | --- | --- | --- | --- |
| Parameter | Estimate | SE | P-value | Estimate | SE | P-value | Estimate | SE | P-value | Estimate | SE | P-value |
| Intercept | 66 | 5.5 | <.0001 | 14 | 1.6 | <.0001 | 18 | 1.7 | <.0001 | 61 | 7.1 | <.0001 |
| Slope | 3.6 | 0.56 | <.0001 | 1.6 | 0.32 | <.0001 | 1.4 | 0.24 | <.0001 | 7.0 | 1.4 | <.0001 |
| Deceleration | 0.026 | 0.0066 | <.0001 | 0.012 | 0.0042 | 0.0022 | 0.012 | 0.0032 | <.0001 | 0.068 | 0.019 | 0.0001 |
| Within-subject covariance | 0.49 | 0.10 | <.0001 | 0.60 | 0.059 | <.0001 | 0.58 | 0.075 | <.0001 | 0.64 | 0.059 | <.0001 |

**Table 11.** Random Effects Covariance Parameter Estimates for Woodcock-Johnson III Standard Scores.

| Parameter | Letter Word ID |  |  | Word Attack |  |  | Passage Comprehension |  |  | Reading Fluency |  |  |
| --- | --- | --- | --- | --- | --- | --- | --- | --- | --- | --- | --- | --- |
|  | Estimate | SE | P-value | Estimate | SE | P-value | Estimate | SE | P-value | Estimate | SE | P-value |
| Intercept | 232 | 19 | <.0001 | 154 | 14 | <.0001 | 189 | 17 | <.0001 | 1121 | 97 | <.0001 |
| Slope | 6.2 | 1.3 | <.0001 | 7.5 | 1.5 | <.0001 | 13 | 2.0 | <.0001 | 175 | 17 | <.0001 |
| Deceleration | 0.019 | 0.014 | 0.08 | 0.048 | 0.019 | 0.0058 | 0.096 | 0.023 | <.0001 | 1.6 | 0.17 | <.0001 |
| Within-subject covariance | 0.58 | 0.070 | <.0001 | 0.51 | 0.11 | <.0001 | 0.68 | 0.044 | <.0001 | 0.86 | 0.014 | <.0001 |

## Discussion

Utilizing a mean-split transformation of a latent phenotype indexing decoding in poor performing students from the longitudinal NHLP, we identified an association between *GARRE1* on chromosome 19 and decoding performance that exceeded the threshold for genome-wide significance, and was well-controlled for ancestry, sex, and SES. We replicated our results in an age-matched subset of the GRaD cohort. In addition, we observed that age moderates the effect of rs2599553 on decoding performance with opposing directions of effect for different quantiles of age.

Further analysis with *Tractor* partitioned a single VCF from admixed subjects into three separate, ancestry-specific files containing alleles from chromosome segments inherited from a single ancestry. This allowed for fine-scale control of population structure in admixed and mixed cohorts, detection of ancestry-specific differences in allele frequencies and effect sizes, and identification of the ancestral source of our primary GWAS signal. eQTL data from the GTEx Project suggested that the GWAS signal likely originated from *GARRE1*, which may play a role in cerebellar development and function. Expression data from Brainspan suggested that there is a developmental change in the expression of *GARRE1*, from equal expression in cortex and cerebellum prenatally, to predominantly cerebellum postnatally through adulthood.

In the NHLP, gene risk group significantly predicted reading skills, as measured by four related dimensions of reading at the beginning of Grade 1. The risk effect was persistent longitudinally across all testing points to the beginning of Grade 5 for all measures. These results indicate that children with the minor allele of rs2599553 begin Grade 1 with lower reading ability and that this gap is maintained throughout the primary school years until at least the beginning of Grade 5. There was no relationship between *GARRE1* risk and the *rate* of acquisition of reading skills.

When the outcome was children’s performance relative to developmental expectations, as represented by the normative sample of the reading test, risk associated with *GARRE1* was also demonstrated. As illustrated by the growth curve analyses (Figure 8a and 8b), these differences in reading performance were maintained at every time point until the end of Grade 5, though there was no relationship between *GARRE1* risk and change in clinical severity over time. Of particular clinical significance, children with one rs2599553 minor allele performed below average and showed the greatest deficits in passage comprehension and reading fluency relative to their peers at the start of Grade 1. By the end of Grade 5, children with the minor allele as a group approached the clinical cutoff for a specific learning deficit in reading.

**Figure 8b.**
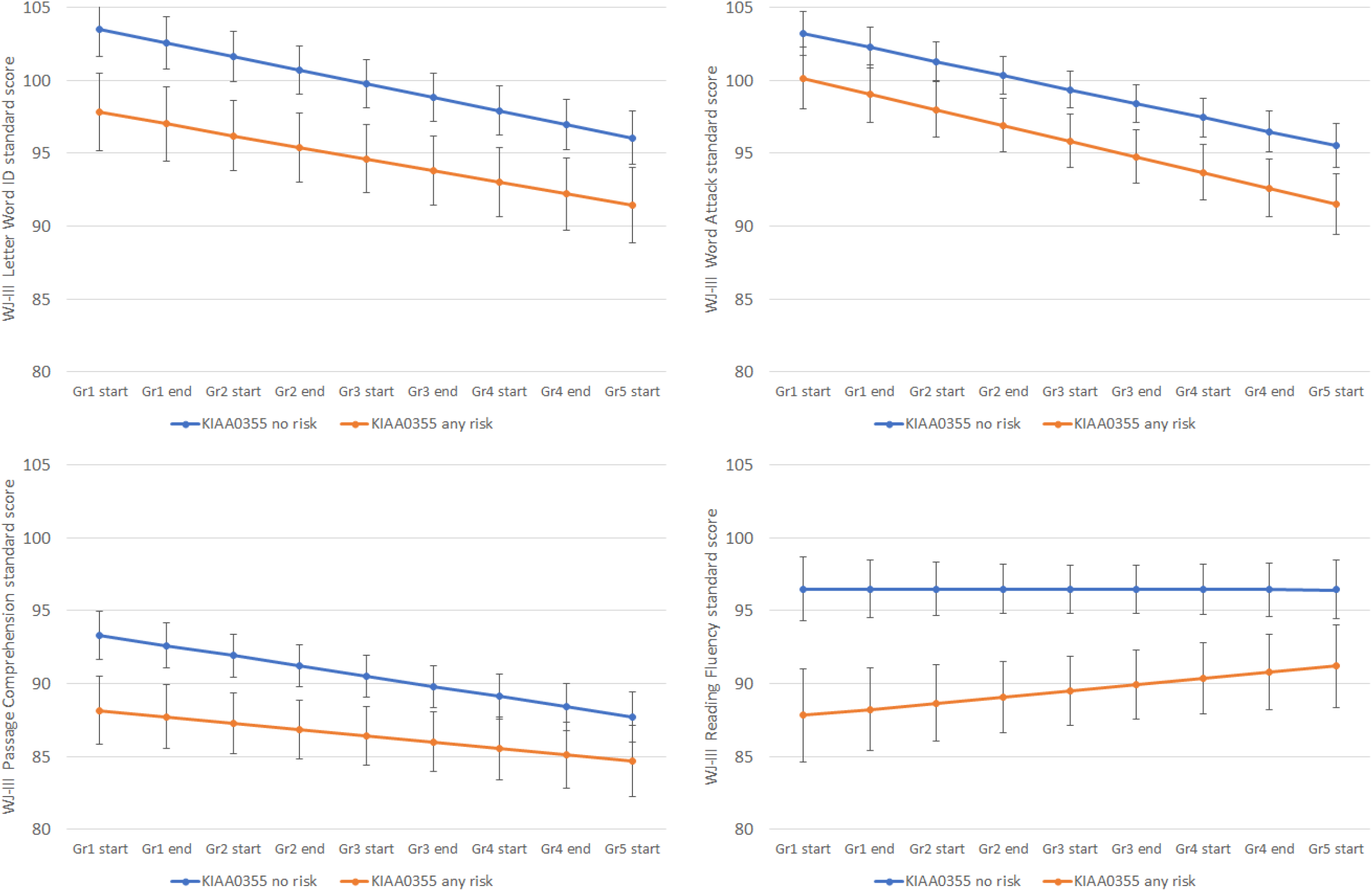
Growth curves depicting model-estimated standard scores on four standardized reading assessments from the Woodcock-Johnson III, and covering the developmental period from the start of Grade 1 until the start of Grade 5. Within each graph, growth curves are presented separately for the two GARRE1 risk groups.

## Limitations

The primary limitation of this study was the small size of the discovery cohort. Underpowered studies have a higher risk of type I errors. We addressed two common sources of type I error in GWAS through chi-squared and ANOVA testing of allele frequency and decoding performance; neither test showed significant deviation for any self-report racial category and the tested source of type I error. While this does not completely exclude the possibility of error, the *post-hoc* statistical testing combined with the robust ancestry correction strengthen the likelihood that the results are valid. Additionally, the general pattern for uncontrolled admixture in GWAS is a global inflation of test statistics across the genome, leading to an elevated genome inflation factor (lambda). The lambda value showed no evidence for significant test statistic inflation. Furthermore, the only locus across the genome to approach statistical significance is the 31 SNP region on chromosome 19.

In addition, referral and recruitment biases amplify as children age. Recruiting young children within the school system can capture a relatively unbiased sample dependent only on schooling rates, but problems in recruiting an unbiased sample of older children with reading problems are amplified by economic (McLaughlin et al., 2014), justice (McGee et al., 1988) and health (Bynner & Parsons, 2001) issues related to RD itself. These dynamics can skew the estimation of gene-behavior relationships.

An interesting question raised by these results is why this locus was never observed in previous studies of reading, especially in light of the effects on development observed in the growth curve analysis. Analysis results from *Tractor* indicate that the association results were driven by African ancestry genetic tracts. To date, only a single GWAS has been published that reported an association with a reading skill that exceeded the threshold for genome-wide significance, and that study had a significant proportion of subjects of non-European ancestry.(Truong et al., 2019) The fact that the signal from the primary GWAS was strongest in AFR tracts in the NHLP, though not unique to them, may partially explain why this locus has remained unreported to date. Most previous studies of reading did not include a significant number of subjects from Hispanic or African American backgrounds, limiting their ability to identify this chromosome 19 locus. Indeed, nearly 80% of all published GWAS are on European cohorts, and this Eurocentricity is even more pronounced when considering only psychiatric and cognitive phenotypes.(Dalvie et al., 2015; Sirugo et al., 2019) The ability of this study to identify a novel signal driven by AFR tracts highlights the utility of including diverse and admixed cohorts in discovering and resolving genetic associations.

### *GARRE1*, *RAC1*, and the Cerebellum

Little is known about the function of *GARRE1* (Granule Associated Rac and RHOG Effector 1; previously known as *KIAA0355*). Using optogenetics, Halil Bagci and coauthors showed physical interaction between GARRE1 and RAC1.(Bagci et al., 2020) In 2018, a proximity mapping experiment localized GARRE1 protein to cytoplasmic granules in HEK293 cells, suggesting a role in mRNA processing and protein expression.(Youn et al., 2018) RAC1 is a relatively well characterized Rho GTPase associated with a diverse collection of cellular processes, including lamellipodia formation. Lamellipodia are transient cell structures associated with cellular migration, including neurons, that have been reported to play a role in reading and language problems.(Guidi et al., 2018) *RAC1* mutations have been associated with severe developmental disorders, including at least one case report associated with cerebellar hypoplasia and microcephaly.(Reijnders et al., 2017)

In neuroimaging research of reading ability, converging evidence has implicated the cerebellum in reading processes (Paulesu et al., 2014). Studies have demonstrated cerebellar activity in a variety of component reading skills including language, phonological, and semantic processing (McDermott et al., 2003; Booth et al., 2007; King et al., 2019). Recent studies of children show differences in cerebro-cerebellar resting state functional connectivity between the cerebellum and left supramarginal gyrus that predict rapid automatized naming (Ang et al., 2020) and differences between children with and without dyslexia on connectivity between the cerebellum and the left inferior frontal gyrus, posterior superior temporal gyrus, and the angular gyrus (Ashburn et al., 2020). Our results lend genetic support to neuroimaging associations between the cerebellum and reading ability. One potential pathway could be that variation in *GARRE1* may lead to modulation of RAC1 activity or expression that presents as subtle changes in the cerebellum connectivity and difficulty with rapid automatized naming. Further studies are needed to understand the role of the cerebellum and *GARRE1* in reading development.

### Reading, Genetics, and Gene-by-Age Effects

Children pass through several developmental windows as they become fluent readers, and as brain circuits mature. This creates changing relationships between genes and behavior and likely underlies the age-dependent effects that we observed in the GRaD sample. The *GARRE1* risk allele had a positive effect in younger children and a negative effect in older children. If other reading-related traits similarly show opposite directions of effect at different ages, a meta-analysis that includes children at a wide range of ages could cancel out and obscure associations. Careful designs that account for gene-by-age effects will be critical in studies that combine datasets to maximize the potential from often underpowered and heterogeneous samples common to the genetics of reading. Prospective studies will ideally be structured cohort or longitudinal designs to capture the changing dynamics over development.

This study highlights the importance of wide and deep phenotyping, longitudinal study design, and inclusion of diverse populations for genetic studies. We demonstrate a viable path for novel genetic discovery and candidate gene identification, even in small primary samples, through the construction of a holistic approach that integrates GWAS, replication, and bioinformatics. Our results add further evidence in support of genetic screening to presymptomatically identify children who are at significant risk for reading deficits. They also suggest that future analyses of the NHLP could show new correlations between genetic variants and variable responses to a comprehensive intervention, a potential clinically useful tool for counseling students and their parents and for modifying curricula.

## Acknowledgements

The authors would like to acknowledge the testers and administrative staff that gathered data for both studies, as well as the subjects and their parents for generously sharing their time and effort as part of this study. This project was supported by the Manton Foundation (A.K.A., M.M.C.D., J.M.B.H., J.C.F., and J.R.G), the Eunice Kennedy Shriver National Institute of Child Health and Human Development (P50HD027802 to J.R.G. and K99HD094902 to D.T.T), and the National Institute of Mental Health (K01MH121659 and T32MH017119 to E.G.A.).

